# An EDS1 EP-domain surface mediating timely transcriptional reprogramming of immunity genes

**DOI:** 10.1101/362921

**Authors:** Deepak D. Bhandari, Dmitry Lapin, Barbara Kracher, Patrick von Born, Jaqueline Bautor, Karsten Niefind, Jane E. Parker

## Abstract

Plant intracellular NLR receptors recognize pathogen interference to trigger immunity. NLR signalling mechanisms have not been resolved. Enhanced disease susceptibility 1 (EDS1) heterodimers are recruited by Toll-interleukin1-receptor domain NLRs (TNLs) to transcriptionally mobilize resistance pathways. Using an Arabidopsis EDS1 heterodimer crystal structure we interrogate the conserved but functionally uncharacterized EDS1 α-helical EP-domain. We identify EP-domain positively charged residues lining a cavity that are essential for TNL immunity signalling, beyond heterodimer formation. Mutating arginine (R493) to alanine creates a weak *EDS1* allele which disables TNL immunity against bacteria producing a virulence factor, coronatine (COR). Arabidopsis plants expressing EDS1^R493A^ are slow to mobilize defence gene expression changes, independently of COR. The transcriptional delay has severe consequences for pathogen resistance and for countering bacterial COR. We uncover a set of host immunity genes whose repression by COR is blocked by wild-type EDS1 but not by EDS1^R493A^ in the TNL response. These data uncover an EDS1 signalling surface lining the heterodimer EP-domain cavity which confers timely transcriptional reprogramming of host defence pathways and blocks bacterial virulence in NLR receptor immunity.

## Introduction

In plants and animals, innate immunity is governed by surface and intracellular receptors. Mammalian innate immune responses provide an initial barrier against microbial infection, and specific pathogen resistance is normally taken over by the adaptive immune system. By contrast, plants depend entirely on panels of germ line-encoded receptors (Jacob *et al*, 2013; Jones *et al*, 2016). Plant recognition of specific pathogen virulence components (known as pathogen effectors) is conferred by intracellular nucleotide-binding/leucine-rich repeat (NLR) receptors in a process called effector-triggered immunity (ETI). NLRs directly or indirectly intercept effectors delivered by pathogens to host cells (Cui *et al*, 2015; Jones *et al*, 2016; Zhang *et al*, 2017). NLR-effector recognition causes receptor conformational activation via an ADP/ATP-dependent switch mechanism, which leads to induction anti-microbial defence pathways, often accompanied by localized host cell death at infection sites (the hypersensitive response, HR).

Two major plant NLR classes are defined principally by their N-terminal domain architectures: those with an N-terminal coiled-coil (CC) domain are known as CNLs (or CC-NLRs) and those with a Toll-interleukin 1 receptor (TIR) domain, as TNLs (or TIR-NLRs) (Jacob et al. 2013; Zhang et al. 2017). A characteristic feature of ETI mediated by the different NLR types is amplification of a similar suite of defence pathways that are mobilized at a lower level by surface pattern-recognition receptors (PRRs) recognizing microbe-associated molecular patterns (MAMPs) in basal immunity (Tao *et al*, 2003; Bartsch *et al*, 2006; Navarro *et al*, 2004; Tsuda *et al*, 2009; Mine *et al*, 2018; Jacob *et al*, 2018). Basal immunity pathways are often targeted by pathogen effectors and the transcriptional reestablishment and bolstering of immunity outputs in ETI is a major driver of resistance (Cui *et al*, 2015; Tsuda & Somssich, 2015). Current evidence suggests that ETI transcriptional reprogramming is facilitated by the early removal of repression of immunity-related components, setting up waves of transcription factor (TF)-orchestrated changes in host cells and tissues (Cui *et al*, 2015; Sun *et al*, 2015; Jacob *et al*, 2018; Zhou *et al*, 2018). Numerous TFs have been found to contribute to ETI governed by CNL and TNL receptors(Cui *et al*, 2015; Birkenbihl *et al*, 2017). Also, several NLRs have nuclear functions (García & Parker, 2009; Inoue *et al*, 2013; Sarris *et al*, 2015; Le Roux *et al*, 2015; Fenyk *et al*, 2015; Cui *et al*, 2015), suggesting that the path between NLR activation and gene expression reprogramming might in some cases be short. However, the mechanisms by which NLRs modulate transcriptional defences are not known.

Another emerging NLR ETI characteristic which distinguishes it from generally weaker basal immune responses, is the contribution of alternative (parallel) transcriptional branches, enabling the plant to compensate for disabling of a particular host resistance sector (Tsuda *et al*, 2009; Kim *et al*, 2014b; Cui *et al*, 2017; Cui *et al*, 2018). ETI buffering of pathways appears to provide robustness against pathogen interference. For example, Arabidopsis ETI conferred by the TNL pair (*RRS1S RPS4*) and a CNL (*RPS2*) receptor recognizing specific effectors delivered by leaf-infecting *Pseudomonas syringae* pv *tomato (Pst*) bacteria, engage a prolonged MAP kinase (MPK3/MPK6) cascade and the nucleocytoplasmic, lipase-like regulator Enhanced Disease Susceptibility1 (EDS1) to protect salicylic acid (SA) responsive gene expression outputs (Tsuda *et al*, 2013; Cui *et al*, 2017). SA is a phenolic hormone produced by the pathogen-induced enzyme, isochorismate synthase1 (ICS1) (Wildermuth *et al*, 2001), and a central component of plant local and systemic immunity against biotrophic pathogens (Fu & Dong 2013). SA accumulation is controlled by an ensemble of TFs operating within a phytohormone network, which enables the plant to prioritize responses to a prevailing stress (Seyfferth & Tsuda, 2014; Pieterse *et al*, 2012).

Different effectors delivered to host cells by fungal, oomycete and bacterial pathogens target SA immunity, often by boosting the SA-antagonising jasmonic acid (JA) hormone system which mediates resistance to necrotrophic pathogens and insects (Kazan & Lyons, 2014; Yang *et al*, 2017). Coronatine (COR) is a potent SA-antagonizing virulence molecule produced by *Pseudomonas* bacteria (including *Pst*) which promotes infection by mimicking plant endogenous bioactive JA-isoleucine (JA-Ile) (Brooks *et al*, 2005; Zheng *et al*, 2012). Like JA-Ile, COR signals by binding to nuclear F-box protein coronatine-insensitive1 (COI1) - jasmonate ZIM-domain (JAZ) coreceptors, which relieves JAZ repression of a basic helix-loop-helix (bHLH) TF, myelocytomatosis oncogene homolog2 (MYC2) (Kazan & Manners, 2013; Zhang *et al*, 2015b). MYC2 is a hub TF for JA, ethylene and abscisic acid signalling, orchestrating numerous stress outputs, including transcriptional dampening of SA accumulation (Kazan & Manners, 2013; Zhang *et al*, 2015b).

In Arabidopsis and other dicotyledenous plant species, *EDS1* is essential for TNL ETI and autoimmune outputs - transcriptional reprogramming, SA accumulation, host cell death and inhibition of pathogen growth (Li *et al*, 2001; Wiermer *et al*, 2005; García & Parker, 2009; Gao *et al*, 2014; Stuttmann *et al*, 2016). Hence, EDS1 is an early convergence point for activated TNLs recognizing different pathogens, and a crucial link to downstream pathways. A small nuclear EDS1 pool is sufficient for Arabidopsis basal immunity against virulent pathogens and TNL ETI, while cytoplasmic EDS1 contributes to pathogen-triggered host cell death (García *et al*, 2010; Rietz *et al*, 2011; Heidrich *et al*, 2011; Stuttmann *et al*, 2016).

To signal in TNL ETI, Arabidopsis EDS1 forms separate heterodimer complexes with each of its sequence-related partners, phytoalexin deficient4 (PAD4) and senescence-associated gene101 (SAG101) (Zhou *et al*, 1998; Jirage *et al*, 1999; Feys *et al*, 2001; Feys *et al*, 2005; Rietz *et al*, 2011; Wagner *et al*, 2013). Analysis of the crystal structure of an EDS1-SAG101 heterodimer, and a structural homology-based model of EDS1-PAD4, showed that the partner N-terminal lipase-like (α/β-hydrolase fold) domains act as a noncatalytic scaffold for interaction and promoting contacts between the partner C-terminal ‘EP’ (from EDS1-PAD4) α-helical bundle domains (Rietz *et al*, 2011; Wagner *et al*, 2013). The heterodimer EP-domains create a cavity. The function of the EP-domains and associated cavity has not been established.

Arabidopsis EDS1 and SA signalling pathways operate as genetically parallel, mutually reinforcing resistance sectors (Tsuda *et al*, 2009; Venugopal *et al*, 2009; Cui *et al*, 2018). We recently reported that EDS1-PAD4 heterodimers, besides promoting *ICS1* expression and SA accumulation, antagonize COR stimulated MYC2 pathways in Arabidopsis TNL ETI against *Pst* bacteria (Cui *et al*, 2018). We established that EDS1-PAD4 antagonism of the COR/JA MYC2-branch is independent of its promotion of the ICS1/SA-branch, leading us to propose a two-pronged EDS1 signalling mechanism in ETI for buffering SA immunity against genetic or pathogen interference (Cui *et al*, 2018).

In this study, we investigate the role of the EDS1 EP-domain in Arabidopsis TNL ETI. By characterizing molecular, transcriptional and disease resistance phenotypes of plants expressing EDS1 structure-guided EP-domain mutations, we identify a positive surface lining the EP-domain heterodimer cavity which is necessary for pathogen resistance and for timely transcriptional reprogramming of immunity genes. Strikingly, the EDS1 EP-domain surface also signals in ETI conferred by a CNL receptor (RPS2), and in both TNL (RRS1 RPS4) and CNL (RPS2) ETI to *Pst* bacteria, EDS1 EP-domain functions can be dissected into three distinct resistance branches.

## Results

### Residues lining the EDS1 heterodimer cavity mediate immunity signalling

The Arabidopsis EDS1-SAG101 heterodimer crystal structure reveals a cavity formed by the partner EP-domains with several conserved positively charged amino acids (lysines and arginines) on the EDS1 and SAG101 surfaces (Fig. S1A, B) (Wagner *et al*, 2013). A similar distribution of positive residues was found in a homology-based structural model of the EDS1-PAD4 heterodimer (Wagner *et al*, 2013). We selected solvent accessible EDS1 lysine (K) and arginine (R) residues that are not part of the heterodimer interface and mutated these individually to alanines (Fig. S1A, C). In a yeast 2-hybrid (Y2H) assay, the EDS1 EP-domain amino acid exchanges did not reduce EDS1-PAD4 interaction compared to the EDS1^LLIF^ lipase-like domain mutant which fails to bind PAD4 (Fig. S1D) (Wagner *et al*, 2013). Constructs of wild-type EDS1 cDNA (cEDS1) and EDS1 EP-domain variants under the *EDS1* native promoter and N-terminally tagged with yellow fluorescent protein (YFP) were transformed into the Arabidopsis Col *eds1-2* null mutant (Wagner *et al*, 2013; Bartsch *et al*, 2006). Primary (T1) transformants for each construct were then inoculated with the oomycete pathogen *Hyaloperonospora arabidopsidis (Hpa*) isolate EMWA1 to test for TNL (*RPP4*) immunity phenotypes, measured against resistant and susceptible controls (Stuttmann *et al*, 2011). Several EP-domain cavity mutants had reduced *RPP4* resistance (Fig. S1C). One mutant, EDS1^R493A^, located in the centre of the cavity (Fig. 1A) was chosen for in-depth analysis because it displayed similarly high *Hpa* EMWA1 disease susceptibility as *eds1-2*. Consistent with the T1 TNL immunity phenotypes, two independent homozygous cEDS1^R493A^ transgenic lines (denoted R493A#1 and #2) were fully susceptible to *Hpa* isolate CALA2 (Fig 1B), recognized in Col by *RPP2A* and *RPP2B TNL* genes (Sinapidou *et al*, 2004), and the bacterial pathogen *Pst AvrRps4* recognized by the nuclear TNL pair RRS1S-RPS4 (Wirthmueller *et al*, 2007; Williams *et al*, 2014; Saucet *et al*, 2015) (Fig. 1C). These data show that EDS1^R493A^ compromises immunity governed by TNLs recognizing an oomycete and bacterial pathogen, without breaking EDS1 heterodimer formation.

**Figure 1.**
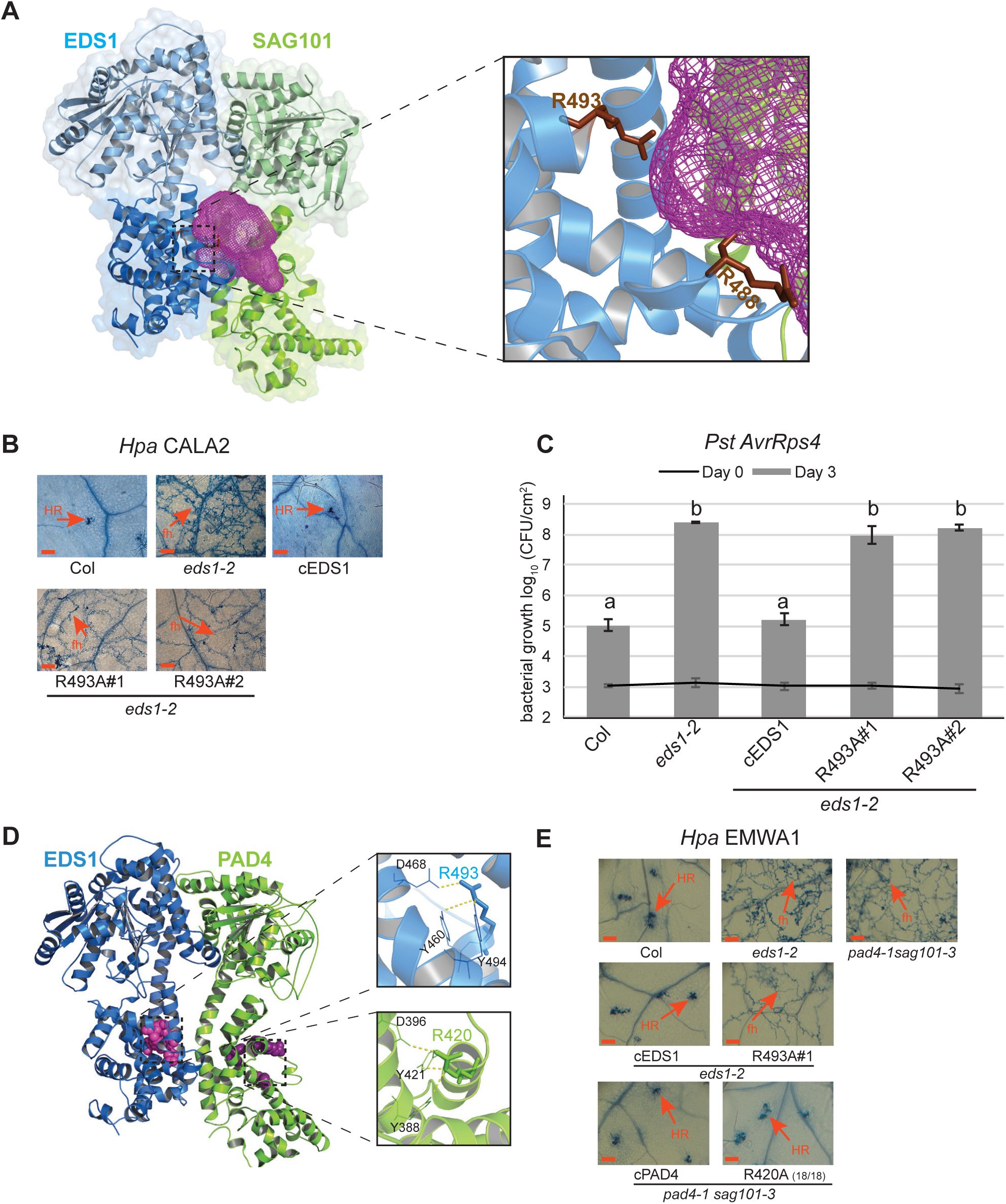
Residues lining the EDS1 heterodimer cavity mediate immunity signalling. **A**. Crystal structure of EDS1 (blue) - SAG101 (green) showing heterodimer formation chiefly driven by the partner lipase-like domains (light tones) and producing a cavity (magenta mesh) formed by the EP-domains. In the zoomout, two conserved EDS1 arginine residues lining the cavity are depicted as sticks (brown). **B**. *RPP2* resistance phenotypes of two-week-old control and transgenic lines expressing YFP-cEDS1 and R493A. *Hpa* EMWA1 infected leaves were stained with trypan blue at 5 dpi. Scale bar represents 100 μm. Images are representative of 24 leaves from two different experiments. HR, hypersensitive response; fh, pathogen free hyphae. **C**. Four-week old Arabidopsis plants of the indicated genotypes were infiltrated with *Pst AvrRps4* (OD_600_ - 0.0005) and bacterial titers determined at 0 and 3 dpi. Bars represent mean of four biological replicates ± SE. Differences between genotypes were determined using ANOVA (Tukey’s HSD, p <0.005). Similar results were obtained in three independent experiments. **D**. Homology model of EDS1 (blue) - PAD4 (green). Conserved residues in the EP-domains of EDS1 (magenta) and PAD4 (purple) are represented as spheres. EDS1 residues line the heterodimer cavity while PAD4 residues are not part of the cavity. The zoomout shows ionic and hydrogen bonds formed by EDS1^R493^ and the equivalent arginine residue in PAD4^R420^ with neighbouring residues. **E**. *RPP4* resistance phenotypes of two-week-old control and homozygous transgenic lines expressing wild-type EDS1, PAD4 and mutated arginine variants. *Hpa* EMWA1 infected leaves were stained with trypan blue at 5 dpi. Scale bar represents 100 μm. The PAD4 R420A image is representative of 18 independent transgenic (T1) plants. HR, hypersensitive response; fh, pathogen free hyphae.

Further examination of the EDS1-SAG101 and EDS1-PAD4 structures revealed a SAG101 and PAD4 EP-domain surface resembling the EDS1^R493A^ patch but facing away from the cavity (Fig. 1D). We mutated PAD4 arginine 420 (PAD4^R420A^) which is predicted to form similar interactions with neighbouring residues as EDS1^R493^ (Fig. 1D), but on the external PAD4 surface. These amino acids are also aligned at the sequence level between EDS1 and PAD4. Wild-type cPAD4 and PAD4^R420A^ controlled by the *PAD4* native promoter and N-terminally tagged with YFP were transformed into Col *pad4-1 sag101-3* mutant in which loss of *SAG101* is compensated for by *PAD4* (Feys *et al*, 2005; Wagner *et al*, 2013). In *RPP4 (Hpa* EMWA1) infection assays of T1 transgenic lines, PAD4 R420A was as resistant as cPAD4 and Col, whereas the EDS1 R493A#1, *eds1-2* and *pad4-1 sag101-3* plants were susceptible (Fig. 1E). These data suggest that the location of positively charged residues, such as EDS1^R493^, within the EP-domain cavity is crucial for TNL immunity.

We tested whether EDS1^R493A^ retains interaction with PAD4 *in planta* by performing transient expression and immunoprecipitation (IP) assays in *eds1-2pad4-1 sag101-3* protoplasts (Cui *et al*, 2018). EDS1^R493A^ fused to a FLAG tag interacted with PAD4-YFP as strongly as wild-type EDS1-FLAG, whereas a non-interacting EDS1^LLIF^-FLAG variant did not bind PAD4-YFP (Fig. S2A). YFP-tagged cEDS1^R493A^ protein in the R493A#1 and R493A#2 transgenic lines had a similar nucleocytoplasmic distribution as YFP-cEDS1 at 24 h post infection (hpi) with *Pst AvrRps4* (Fig. S2B), indicating that loss of TNL immunity is not due to failed EDS1 nuclear accumulation (García *et al*, 2010; Stuttmann *et al*, 2016). However, confocal microscopy imaging of multiple samples showed that YFP-cEDS1^R493A^ nucleocytoplasmic fluorescence was lower than YFP-cEDS1 (Fig. S2B). On a protein blot probed with α-GFP antibodies, cEDS1^R493A^-YFP accumulation in R493A#1 was indeed less than cEDS1-YFP in mock‐ and *Pst* AvrRps4-inoculated leaf extracts (Fig. S2D). To test whether the EDS1^R493A^ defect in TNL immunity is due to its low accumulation, we generated transgenic *eds1-2* lines expressing genomic EDS1-YFP or EDS1^R493A^-YFP (denoted gEDS1 and gR493A) because genomic EDS1 is generally more highly expressed than cEDS1 (Stuttmann *et al*, 2016). Although gEDS1 and gR493A accumulated to higher levels than cEDS1 in mock-treated tissues and to similar levels after *Pst AvrRps4* infection (Fig. S2D), the gR493A transgenic line was as susceptible to *Pst AvrRps4* as R493A#1 or *eds1-2*, compared to Col, gEDS1 and cEDS1 plants (Fig. S2C). These data show that impaired TNL (*RRS1S-RPS4*) resistance in R493A is not a consequence of low protein accumulation. We concluded that EDS1^R493^ lining the EP-domain cavity confers an important signalling property on the EDS1-PAD4 heterodimer.

### EDS1^R493A^ delays TNL transcriptional reprogramming

TNL/EDS1 bacterial immunity divides into two mutually reinforcing resistance branches: one involving SA synthesized by EDS1-induced *ICS1*, the other working independently of SA and involving EDS1 antagonism of MYC2 (Cui *et al*, 2017; Cui *et al*, 2018). To check if SA accumulation is affected in R493A, we measured free (active) SA accumulation in leaves of Col, *eds1-2* and cEDS1 and R493A plants at 0, 8 and 24 hpi with *Pst AvrRps4*. At 8 hpi, SA levels in R493A lines #1 and #2 remained low, resembling the *eds1-2* null mutant (Fig. 2A). At 24 hpi, the R493A lines but not *eds1-2* recovered SA accumulation to similar levels as cEDS1 and Col (Fig. 2A). Low SA accumulation at 8 hpi correlated with reduced expression of the SA-marker gene *PR1* (*pathogenesis related 1*) in R493A compared to cEDS1 and Col at 24 hpi (Fig. 2B). These data indicate that EDS1^R493A^ is not a complete loss-of-function mutation but instead slow to mobilize the SA branch of TNL immunity. Therefore, one function of the EDS1 EP-domain, which is compromised in R493A, is to promote the SA immunity branch.

**Figure 2.**
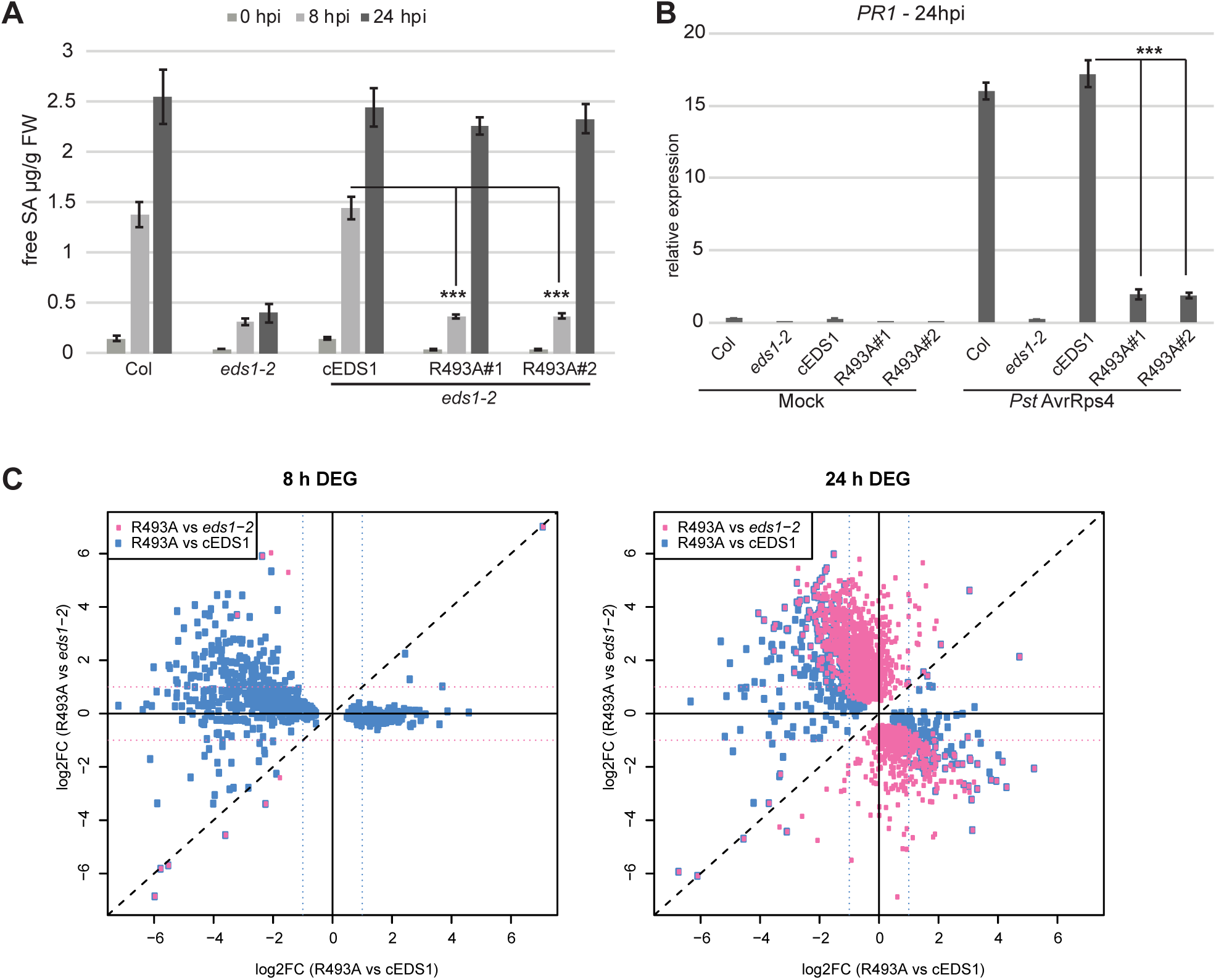
EDS1^R493A^ delays TNL transcriptional reprogramming. **A**. Four-week-old plants were infiltrated with *Pst AvrRps4* (OD_600_ – 0.005) and free SA was quantified at 0, 8 and 24 hpi. Bars represent means ± SE of four biological replicates. Differences between genotypes were analysed using t-test (Bonferroni corrected, p<0.05). Similar results were obtained in two independent experiments. **B**. Four-week-old plants were infiltrated with 10mM MgCh (mock) or *Pst AvrRps4* (OD_600_ – 0.005), and leaf samples collected at 24 hpi. *PR1* transcripts were measured using qRT-PCR and normalized to *GapDH*. Bars represent means ± SE of 3 biological replicates. Difference between genotypes were analysed using t-test (Bonferroni corrected, p<0.05). **C**. A 2-dimensional scatter plot comparing R493A vs *eds1-2* and R493A vs cEDS1 at 8 and 24 hpi with *Pst AvrRps4*. Plots depict differentially expressed genes (DEG) between R493A vs *eds1-2* (pink dots) and R493A vs cEDS1 (blue dots). DEG represented were filtered with a |log2 FC|≥1, FDR≤0.05.

EDS1 SA-independent signalling operating in parallel with the EDS1-promoted SA/ICS1 branch enables the plant to buffer against a disabled SA sector in ETI (Tsuda *et al*, 2009; Venugopal *et al*, 2009; Cui *et al*, 2018). We performed RNA-seq to interrogate the EDS1^R493A^ defect in TNL (*RRS1S RPS4*) transcriptional reprogramming. Four-week-old Col, *eds1-2*, cEDS1 and R493A (line #1) plants were infiltrated with *Pst AvrRps4* and three independent biological replicates for each line were processed and analysed at 0, 8 and 24 hpi (see Methods). In a 2-D scatter plot, at 8 hpi there were only 12 DEGs between R493A and *eds1-2* (Fig. 2C (pink boxes) and Table S1). These included *EDS1* (log_2_ FC 6.03, FDR=9.92E-03) and *PBS3* (log_2_ FC 3.7, FDR=2.65E-03). Therefore, at the level of TNL-triggered SA accumulation and transcriptional reprogramming, R493A behaves like the *eds1-2* null mutant at 8 hpi (Fig. 2 A, C). At 24 hpi, R493A had a markedly different expression profile to *eds1-2* (Fig. 2C) with 2053 DEG (|log_2_ FC|≥1, FDR<0.05). R493A did not fully recover at 24 hpi as there were 576 DEG ((|log_2_ FC|≥1, FDR<0.05) between R493A and cEDS1 compared with 5993 DEG ((|log_2_ FC|≥1, FDR<0.05) between *eds1-2* and cEDS1 (Fig. 2C). These gene expression profiles show that EDS1^R493A^ fails to mobilize TNL transcriptional reprogramming at 8 hpi but recovers largely at 24 hpi. Together with the disease susceptibility phenotypes of R493A lines #1 and #2 (Fig. 1B, C), we concluded that recovery of gene expression in R493A at 24 hpi is too late to halt pathogen growth. Hence, integrity of the EDS1 EP-domain appears to be critical for timely TNL-triggered transcriptional defence reprogramming.

### EDS1^R493A^ fails to antagonize bacterial COR stimulated MYC2 in ETI

*Pst* DC3000 produces the JA-Isoleucine (JA-Ile) mimic coronatine (COR) to promote JA response pathways and suppress SA signalling (Brooks *et al*, 2005; Zheng *et al*, 2012; Geng *et al*, 2014). One function of the EDS1-PAD4 heterodimer in *RRS1S RPS4* ETI is to inhibit COR stimulation of MYC2-regulated JA pathway genes independently of *ICS1* (Cui *et al*, 2018).

Because R493A displayed a general delay in gene expression reprogramming (Fig. 2C) and accumulation of SA (Fig. 2A), we tested whether EDS1^R493A^ is defective in antagonizing COR promoted bacterial infection. For this, *Pst AvrRps4* or the COR-deficient *Pst* Δ*cor AvrRps4* strain were infiltrated into leaves of Col, *eds1-2*, cEDS1 and R493A lines #1 and #2, and bacterial growth measured at 3 dpi. As expected, Col and cEDS1 were equally resistant to *Pst AvrRps4* and *Pst* Δ*cor AvrRps4*, reflecting strong EDS1 antagonism of COR stimulated bacterial growth in *RRS1S RPS4* ETI (Fig. 3A). The *eds1-2* mutant was susceptible to both *Pst AvrRps4* strains but supported 1.5-log lower *Pst* Δ*cor AvrRps4* growth compared to *Pst AvrRps4*, indicating both COR stimulated and COR-independent bacterial growth effects in the absence of *EDS1* (Fig. 3A) (Cui *et al*, 2018). Notably, while R493A lines displayed *eds1-2* level susceptibility to *Pst AvrRps4*, they were as resistant as Col and cEDS1 to *Pst* Δ*cor AvrRps4* (Fig. 3A). Therefore, loss of R493A resistance to *Pst AvrRps4* is due to its failure to antagonize bacterial COR, presumably signalling via MYC2. To verify this, we crossed cEDS1 and R493A#1 into an *eds1-2 myc2-3* mutant background. Whereas R493A (in *eds1-2*) was as susceptible as *eds1-2*, R493A *eds1-2 myc2-3* was as resistant as Col and cEDS1 (Fig. 3B), indicating that EDS1^R493A^ recovers resistance to *Pst AvrRps4* when *MYC2* is mutated. These data point to a defect of EDS1^R493A^ in counteracting COR-dependent, MYC2 promoted bacterial growth in TNL ETI. With respect to *Pst AvrRps4*, therefore, EDS1^R493A^ loss of resistance is conditional on COR stimulation of the MYC2 JA signalling branch.

**Figure 3.**
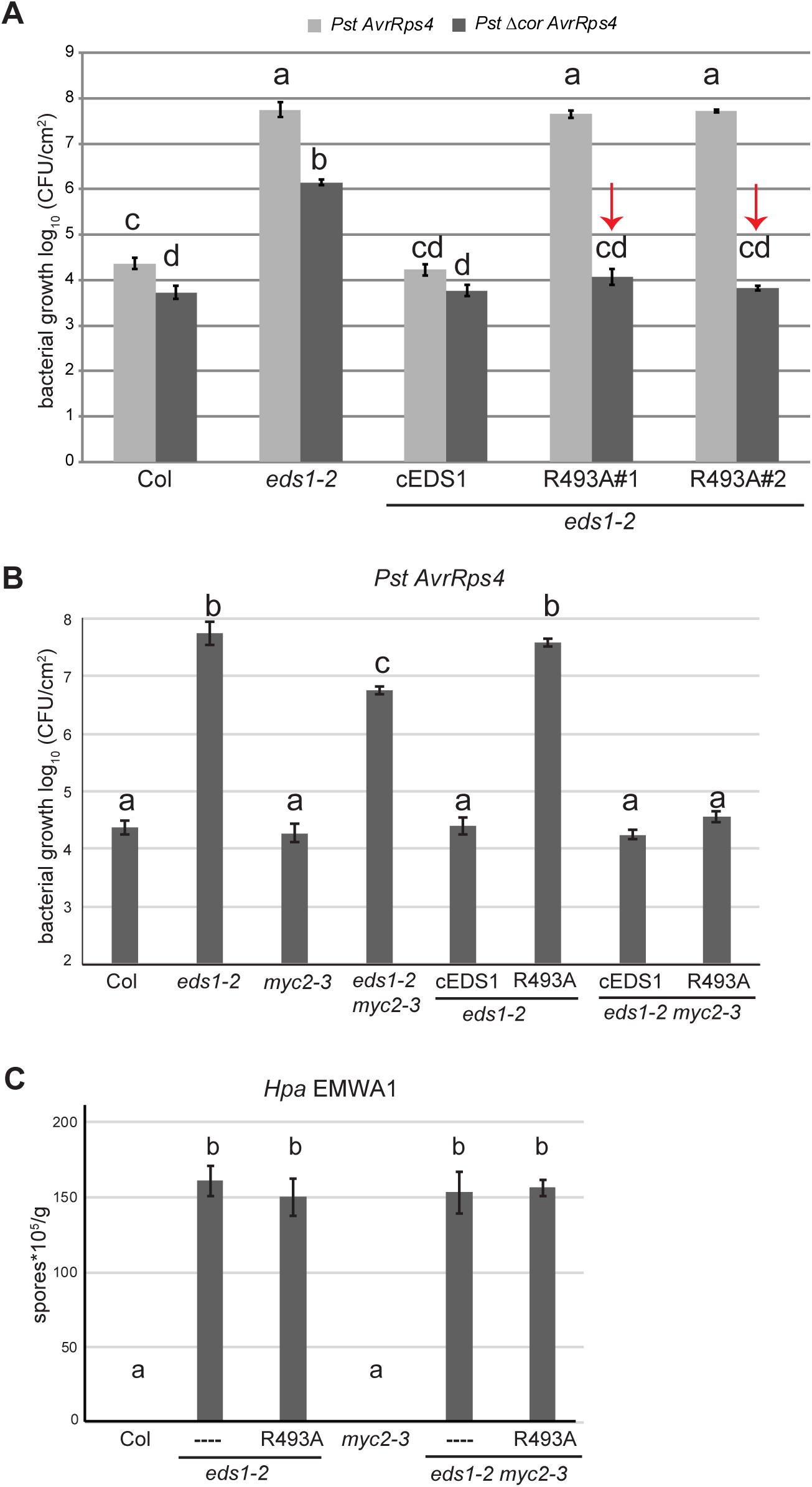
EDS1^R493A^ fails to antagonize bacterial COR-stimulated MYC2 in TNL ETI. **A**. Four-week-old Arabidopsis plants of the indicated genotypes were infiltrated with *Pst AvrRps4* or *Pst* Δ*cor AvrRps4* (OD_600_ - 0.0005). Bacterial titers were determined at 0 and 3 dpi. No significant difference was observed between lines and treatments at 0 dpi. Bars represent mean of four biological replicates ± SE. Differences between genotypes were analysed using ANOVA (Tukey’s HSD, p-value <0.005). Similar results were obtained in three independent experiments. **B**. Four-week-old Arabidopsis plants of the indicated genotypes were infiltrated with *Pst AvrRps4* (OD_600_ - 0.0005). Bacterial titers were determined at 0 and 3 dpi. No significant differences were observed at 0 dpi. Bars represent mean of four biological replicates ± SE. Differences between genotypes were analysed using ANOVA (Tukey’s HSD, p-value <0.005). Similar results were obtained in three independent experiments. **C**. The indicated genotypes were infected with *Hpa* EMWA1 and oomycete sporulation was quantified at 5 dpi. Data from three independent experiments were combined and differences between genotypes analysed using ANOVA (Tukey’s HSD, p<0.005).

Because EDS1^R493A^ was also compromised in TNL (*RPP4* and *RPP2A, B*) immunity to *Hpa* (Fig. 1B, E), we tested whether there is a recovery of *RPP4* resistance to *Hpa* EMWA1 in the *eds1-2 myc2-3* transgenic lines. Col and *myc2-3* expressed full *RPP4* resistance after quantifying EMWA1 sporulation on leaves (Fig. 3C). There were similar high levels of EMWA1 sporulation in *eds1-2*, *eds1-2 myc2-3*, R493A *eds1-2* and R493A *eds1-2 myc2-3* (Fig. 3C). These data show that increased susceptibility to *Hpa* EMWA1 in R493A is not dependent on *MYC2*. Therefore, the EDS1 EP-domain and associated heterodimer cavity have broader functions in TNL immunity than antagonizing MYC2.

### EDS1^R493A^ TNL signalling delay is compounded by bacterial COR

Because COR-activated MYC2 represses SA accumulation (Brooks *et al*, 2005; Zheng *et al*, 2012; Cui *et al*, 2018) and EDS1^R493A^ causes delay in SA accumulation at 8 hpi with *Pst AvrRps4* (Fig. 2A), we tested whether resistance of R493A to *Pst* Δ*cor AvrRps4* (Fig. 3A) is due to restored SA. Col and cEDS1 accumulated similar levels of free SA at 8 hpi with *Pst AvrRps4* and higher SA in response to *Pst* Δ*cor AvrRps4*, consistent with COR dampening of SA accumulation (Fig. 4A). The *eds1-2* null mutant failed to accumulate SA in response to either strain (Fig. 4A), fitting with EDS1 promotion of *ICS1* expression and SA independently of its suppression of MYC2 in TNL (*RRS1S RPS4*) immunity (Cui *et al*, 2018). Strikingly, R493A lines #1 and #2 failed to accumulate SA in response to *Pst AvrRps4* and *Pst* Δ*cor AvrRps4* (Fig. 4A). Therefore, delayed SA accumulation in R493A is not caused by its failure to antagonize bacterial COR. These data suggest that antagonism of COR/MYC2 signalling and promotion of SA accumulation are distinct properties of the EDS1-PAD4 heterodimer EP-domain, and more precisely EDS1^R493^ lining the EP cavity, in TNL ETI.

**Figure 4.**
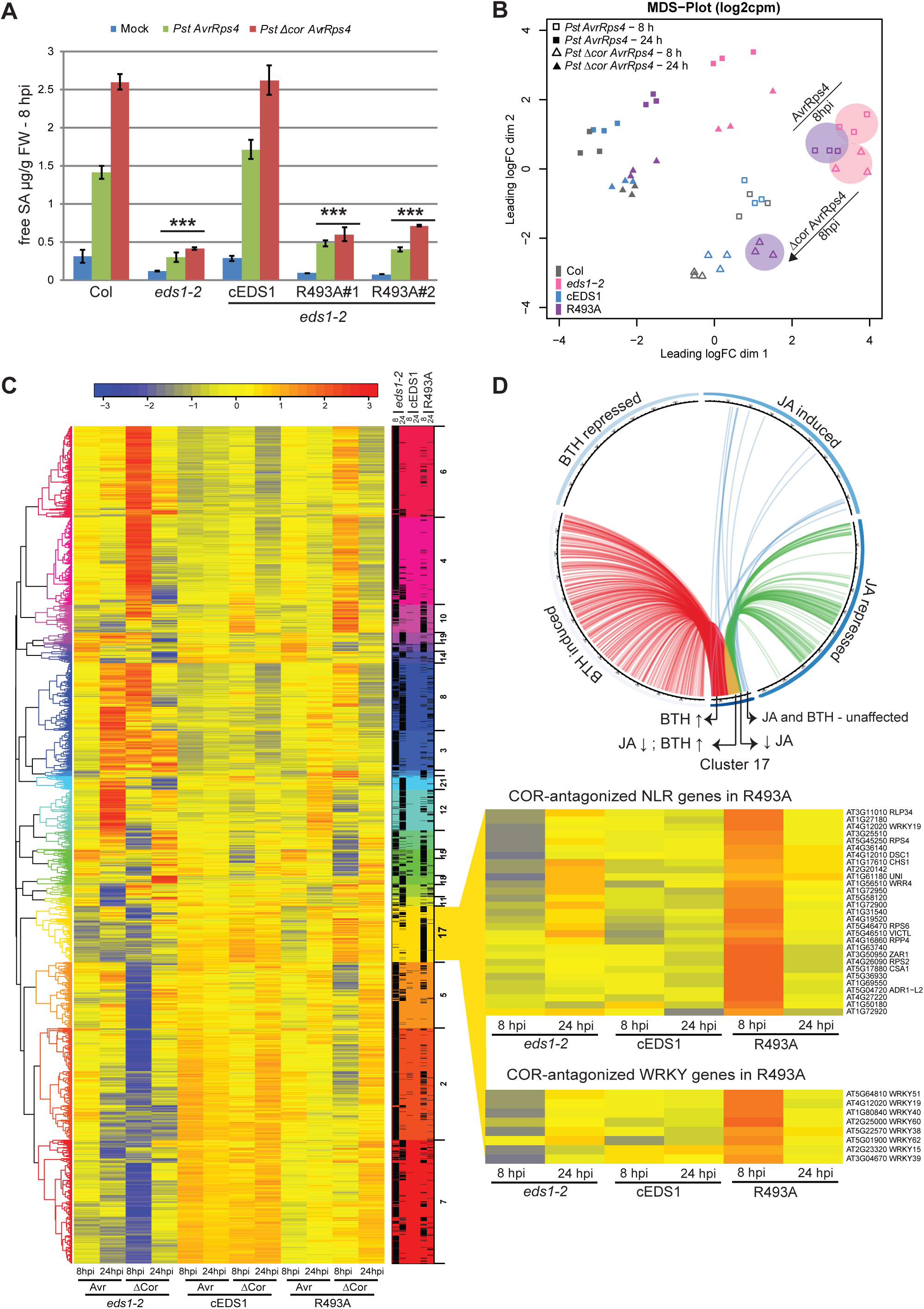
Delayed immune signalling in R493A mutant plants is independent of COR. **A**. Four-week-old plants were infiltrated with 10mM MgCl2 (mock), *Pst AvrRps4* or *Pst* Δ*cor AvrRps4* and free SA was quantified at 8 hpi. Bars represent means ± SE of three biological replicates. Difference between genotypes were analysed using t-test (Bonferroni corrected, p<0.05). Similar results were observed in three independent experiments. **B**. A multidimension scaling (MDS) plot of differentially expressed genes showing R493A transcriptional changes at 8 (open symbols) and 24 hpi (closed symbols). Encircled samples of *eds1-2* (pink) and R493A (purple) highlight transcriptional trends with and without bacterial COR in *Pst AvrRps4*-triggered immunity. **C**. A heatmap depicting DEG at 8 and 24 hpi normalized to Col (p<0.05) after hierarchical clustering. Samples were harvested at 8 and 24 hpi with *Pst AvrRps4* (Avr) and *Pst* Δ*cor AvrRps4* (ऴCor). Cluster #17 contains genes that are upregulated at 8 hpi with *Pst* Δ*cor AvrRps4* but not *Pst AvrRps4* in R493A only. Expansion (right) highlights a subset of *NLR* and *WRKY*genes in cluster #17 that are involved in immunity whose expression is antagonized by COR in R493A (see Table S3). **D**. Circos plot showing overlap of 383 genes in cluster #17 with genes regulated by JA and BTH from other studies (Hickmann et al., 2017; Yang et al., 2017). Genes differentially expressed in cluster #17 and other datasets are marked by connecting lines. Genes repressed by JA and induced by BTH (red and green lines converging to yellow, 113); genes repressed by JA and not expressed by BTH (green lines, 19); genes induced by BTH and not expressed in JA dataset (red lines, 169).

Next, we examined which gene expression sectors affected by bacterial COR might explain the compromised TNL ETI in R493A, by performing an RNA-seq analysis at 8 and 24 hpi with *Pst* Δ*cor AvrRps4*. To compare between *Pst AvrRps4* and *Pst* Δ*cor AvrRps4* RNA-seq experiments, gene expression data for all lines was normalized against the respective Col control within each treatment (see Methods). A multi-dimensional scaling (MDS) plot shows that cEDS1 and Col expression profiles clustered together at 8 and 24 hpi for both treatments (Fig. 4B). At 8hpi, R493A clustered away from cEDS1 and *eds1-2* with *Pst* Δ*cor AvrRps4* but close to *eds1-2* with *Pst AvrRps4* (Fig. 4B, coloured circles). At 24hpi, the R493A transcriptome was similar to cEDS1 with *Pst* Δ*cor AvrRps4* (filled triangles) but distinct from cEDS1 with *Pst AvrRps4* (filled squares) (Fig. 4B). This analysis shows that EDS1^R493A^ causes a general delay in TNL gene expression reprogramming which is exacerbated by bacterial COR. The trend fits with the SA accumulation profiles (Fig. 4A) and emphasizes the importance of EDS1 EP-domain for rapid transcriptional mobilization of multiple pathways besides blocking bacterial COR actions in TNL ETI. We compared our transcriptome data at 8 hpi with publicly available SA‐ and JA-responsive transcriptomes (Tsuda *et al*, 2009; Hickman *et al*, 2017). This showed that R493A exhibits lower expression of genes regulated by *ICS1* compared to cEDS1 regardless of bacterial COR status (Fig. S3A). In contrast, genes that are repressed by JA were also reduced in R493A infiltrated with *Pst AvrRps4* but not with *Pst* Δ*cor AvrRps4* (Fig. S3A). Thus, the R493A transcriptome reflects both slow mobilisation of SA pathways and a defect in antagonising COR/MYC2-regulated JA pathways.

We next searched for DEG between *Pst AvrRps4* and *Pst* Δ*cor AvrRps4* treatments in the different lines at 8 and 24 hpi. After hierarchical clustering, a derived expression heatmap revealed the extent of delay in R493A at 8 hpi compared to cEDS1, both in the presence and absence of bacterial COR (Fig. 4C). Only 75 (8 hpi) and 3 (24 hpi) DEG spread across different clusters were found for cEDS1, reinforcing the notion that EDS1 effectively antagonizes COR in TNL ETI. There were major expression differences between *Pst AvrRps4* and *Pst* Δ*cor AvrRps4* treatments (2810 at 8 hpi and 987 at 24 hpi) in the *eds1-2* null mutant, consistent with EDS1 loss-of-function failing to counter COR effects. While R493A was slow in ETI transcriptional reprogramming against both *Pst AvrRps4* and *Pst* Δ*cor AvrRps4* (Fig. 4B), there remained 1009 (8 hpi) and 210 (24 hpi) DEG (|log_2_ FC|≥1, FDR≤0.05) between these treatments in R493A (Table S2), indicating that the transcriptional delay in R493A is compounded by COR. The 210 DEG found in R493A at 24 hpi mostly overlapped with DEG in *eds1-2*. These expression changes are therefore probably not important for TNL immunity to *Pst AvrRps4*. Because R493A is susceptible to *Pst AvrRps4* but resistant to *Pst* Δ*cor AvrRps4* infection (Fig. 3A), we reasoned that EDS1 interference with COR-dependent expression changes before or at 8 hpi is crucial for halting bacterial growth in the TNL immune response.

### COR represses a distinct set of immunity-related genes in R493A

One expression cluster (#17) stood out in the above analysis because it contains EDS1-dependent DEG at 8 hpi that are more highly expressed in R493A with *Pst* Δ*cor AvrRps4* compared to *Pst AvrRps4* (Fig. 4C, Table S3), suggesting there is targeted repression of a set of genes by COR in R493A but not in Col or cEDS1 lines (which effectively counter COR effects) in TNL immunity. Cluster #17 comprises 383 genes associated with Gene Ontology (GO) terms phosphorylation (37/376 eg. *MPK3*, *MPK2*), cell death (18/376 eg. *RPS2*, *NPR1*, *CPR5*) and defence response (73/376 eg. *RPS4*, *RPP4*, *ADR1-L2*) (Table S4). Notable members of cluster #17 are functionally defined *NLR* (*TNL* and *CNL*) and *WRKY* family TF genes (Fig. 4C, Table S3).

We found a significant overlap between cluster #17 genes and genes regulated by the SA analogue BTH (Yang *et al*, 2017) or JA (Hickman *et al*, 2017). Here, cluster #17 genes fall into four categories (Fig. 4D; Table S5), the largest of which (i) has genes induced by BTH and not repressed by JA (169/383), followed by (ii) genes induced by BTH and repressed by JA (113/383), (iii) genes not responsive to either BTH or JA (82/383), and a small group (iv) genes that are repressed by JA only (19/383). This comparison suggests that TNL/EDS1 signalling at 8 hpi protects a large number of SA/JA responsive but also SA/JA-unrelated immunity genes from bacterial COR repression. Only 30% (118/383) cluster #17 genes overlap with a set of ETI-associated genes that were extracted from transcriptomic analyses of the Arabidopsis *RPS2* (CNL) response to AvrRpt2 expressed *in planta* or delivered by *Pst* bacteria (Hatsugai *et al*, 2017; Mine *et al*, 2018) (Fig. S3B). Therefore, the cluster #17 represents a new and potentially interesting ETI-related gene set. We did not detect significant enrichment of particular cis-regulatory elements in the promoters of cluster #17 genes (by MEME, http://meme-suite.org), suggesting that these genes are controlled by multiple TFs.

In summary, comparison of cEDS1, *eds1-2* and R493A transcriptomes between *Pst AvrRps4* and *Pst* Δ*cor AvrRps4* treatments uncovered a set of TNL/EDS1-controlled genes (cluster #17) whose repression in response to bacterial COR at 8 hpi is inadequately blocked by EDS1^R493A^. Hence, R493A reveals a function of the EDS1 heterodimer EP-domain in countering bacterial COR repression of host immunity genes within the first 8 h of TNL ETI.

### A positive charge at EDS1^R493^ is essential for TNL immunity

We have shown that mutations of positively charged residues in the EDS1 EP cavity impaired EDS1 TNL functions (Fig. S1C, 1B, 1C). To test the importance of the charge at R493, we generated positively (lysine, EDS1^R493K^) and negatively (glutamate, EDS1^R493E^) charged variants of EDS1^R493^. Like EDS1^R493A^, YFP-tagged EDS1^R493K^ and EDS1^R493E^ displayed wild-type nucleocytoplasmic localization in transient expression assays (Fig. S4A). The FLAG-tagged EDS1^R493^ variants interacted with PAD4-YFP in IP experiments (Fig. S4B), consistent with the charge at the EP cavity not affecting heterodimer formation. Additionally, none of the FLAG-tagged EDS1^R493^ variants altered interactions between PAD4-YFP and StrepII-HA (SH)-tagged MYC2 in IPs of transiently expressed proteins (Fig. S4C), indicating that an altered EDS1^R493^ charge does not disturb EDS1-PAD4 association with MYC2.

Two independent homozygous EDS1^R493K^ and EDS1^R493E^ transgenic lines (respectively, R493K and R493E#1 and #2) in *eds1-2* were tested for TNL (*RPP4*) resistance to *Hpa* EMWA1. R493E was as susceptible as *eds1-2* and R493A, whereas R493K expressed full *RPP4* resistance, as monitored by trypan blue staining of infected leaves (Fig. 5A). Therefore, a positive charge at EDS1 amino acid 493 rather than an arginine *per se* is required for TNL immunity to *Hpa* EMWA1.

**Figure 5.**
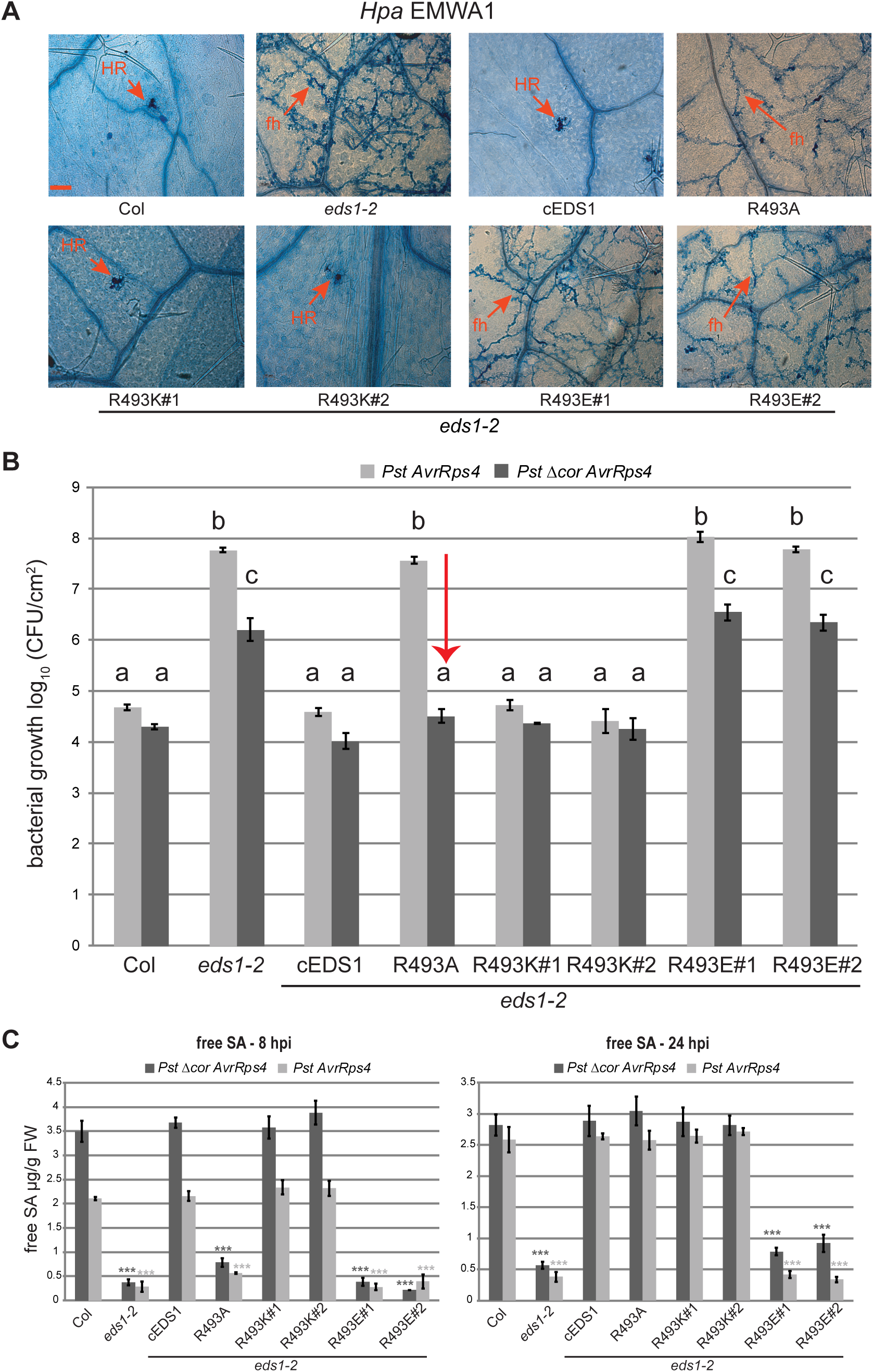
A positive charge at EDS1^R493^ is essential for TNL immunity. **A**. *RPP2* resistance phenotypes of two-week-old control and Arabidopsis transgenic lines expressing cEDS1 and R493 mutants, as indicated. *Hpa* EMWA1 infected leaves were stained with trypan blue at 5 dpi. Scale bar represents 100 μm. Each image is representative of >18 leaves from two independent experiments. HR, hypersensitive response; fh, pathogen free hyphae. **B**. Four-week-old Arabidopsis plants of the indicated genotypes were infiltrated with *Pst AvrRps4* and *Pst* Δ*cor AvrRps4* (OD_600_ - 0.0005). Bacterial titers were determined at 3 dpi. Bars represent means of three biological replicates ± SE. Differences between genotypes were analysed using ANOVA (Tukey’s HSD, p<0.05). Similar results were obtained in three independent experiments. C. Four-week-old plants were infiltrated with *Pst AvrRps4* or *Pst* Δ*cor AvrRps4* and free SA was quantified at 8 and 24 hpi. Bars represent means ± SE of three biological replicates. Differences between genotypes within treatment were analysed using Student’s t-test (Bonferroni corrected, *** - p<0.05) relative to Col. Similar results were obtained in two independent experiments.

Because R493A susceptibility to *Pst AvrRps4* was conditional on COR (Fig. 3A), we tested the TNL (*RRS1S RPS4*) responses of R493K and R493E lines to *Pst AvrRps4* and *Pst* Δ*cor AvrRps4*. Measured against Col, cEDS1 and *eds1-2*, R493K was fully resistant while R493E was fully susceptible to both *Pst AvrRps4* strains, whereas R493A was susceptible only to *Pst AvrRps4* (Fig. 5B). Free SA accumulation in the R493K and R493E lines at 8 and 24 hpi with *Pst AvrRps4* or *Pst* Δ*cor AvrRps4* mirrored, respectively, cEDS1 and *eds1-2* (Fig. 5C). The same trend was found at the level of expression of marker genes (*EDS1*, *PAD4* and *ICS1*) for the EDS1/PAD4-induced SA immunity branch at 8 hpi (Fig. S5A). These SA accumulation, gene expression and disease resistance phenotypes show that R493K phenocopies wild-type and R493E the *eds1-2* null mutant. We performed qRT-PCR analysis of MYC2-branch JA response marker genes *SA methyl transferase 1* (*BSMT1*), *JAZ10* and *Vegetative Storage Protein* 1 (*VSP1*) in the R493 variants at 24 hpi with *Pst AvrRps4* (Cui *et al*, 2018). Here, R493K behaved like wild-type cEDS1 and R493E like *eds1-2* (Fig. S5B). By contrast, R493A repressed *BSMT1, JAZ10* and *VSP1* MYC2-branch genes almost as strongly as cEDS1 (or Col) (Fig. S5B). Hence, EDS1^R493A^ antagonized the COR-stimulated MYC2-branch at 24 hpi but was unable to counter *Pst AvrRps4* infection in ETI (Fig. 3A). Therefore, near wild-type suppression of MYC2-branch genes in R493A plants at 24 hpi was insufficient for TNL ETI. Together, these data show that a positive charge at EDS1^R493^ is critical for timely defence gene expression changes and pathogen resistance in TNL ETI.

### EDS1^R493A^ residual signalling function is specific to ETI

EDS1-PAD4 complexes confer basal immunity to virulent *Pst* DC3000 in the absence of TNL-effector recognition (Jirage *et al*, 1999; Feys *et al*, 2001; Rietz *et al*, 2011; Wagner *et al*, 2013). We therefore tested whether EDS1^R493A^ restriction of *Pst* DC3000 growth is conditional on bacterial COR by infiltrating leaves of the different EDS1^R493^ transgenic lines with *Pst* DC3000 or *Pst* Δ*cor*. Col, cEDS1 and R493K expressed similar resistance to each bacterial strain, with ~ 10-fold higher *Pst* DC3000 growth than *Pst* Δ*cor* at 3 dpi (Fig. 6A). The *eds1-2*, R493A and R493E lines were similarly susceptible to each strain, with ~ 100-fold higher *Pst* DC3000 growth compared to *Pst* Δ*cor* (Fig. 6A). Therefore, R493A does not recover basal resistance to *Pst* Δ*cor*, whereas it recovers TNL (*RRS1SRPS4*) ETI to *Pst* Δ*cor AvrRps4* (Fig. 3A, 5B). This result shows that while a positive charge at EDS1^R493^ is necessary for TNL ETI and basal immunity against *Pst* bacteria, EDS1^R493A^ is partially equipped for TNL ETI but unequipped for basal immunity even without resistance repressive effects of bacterial COR.

**Figure 6.**
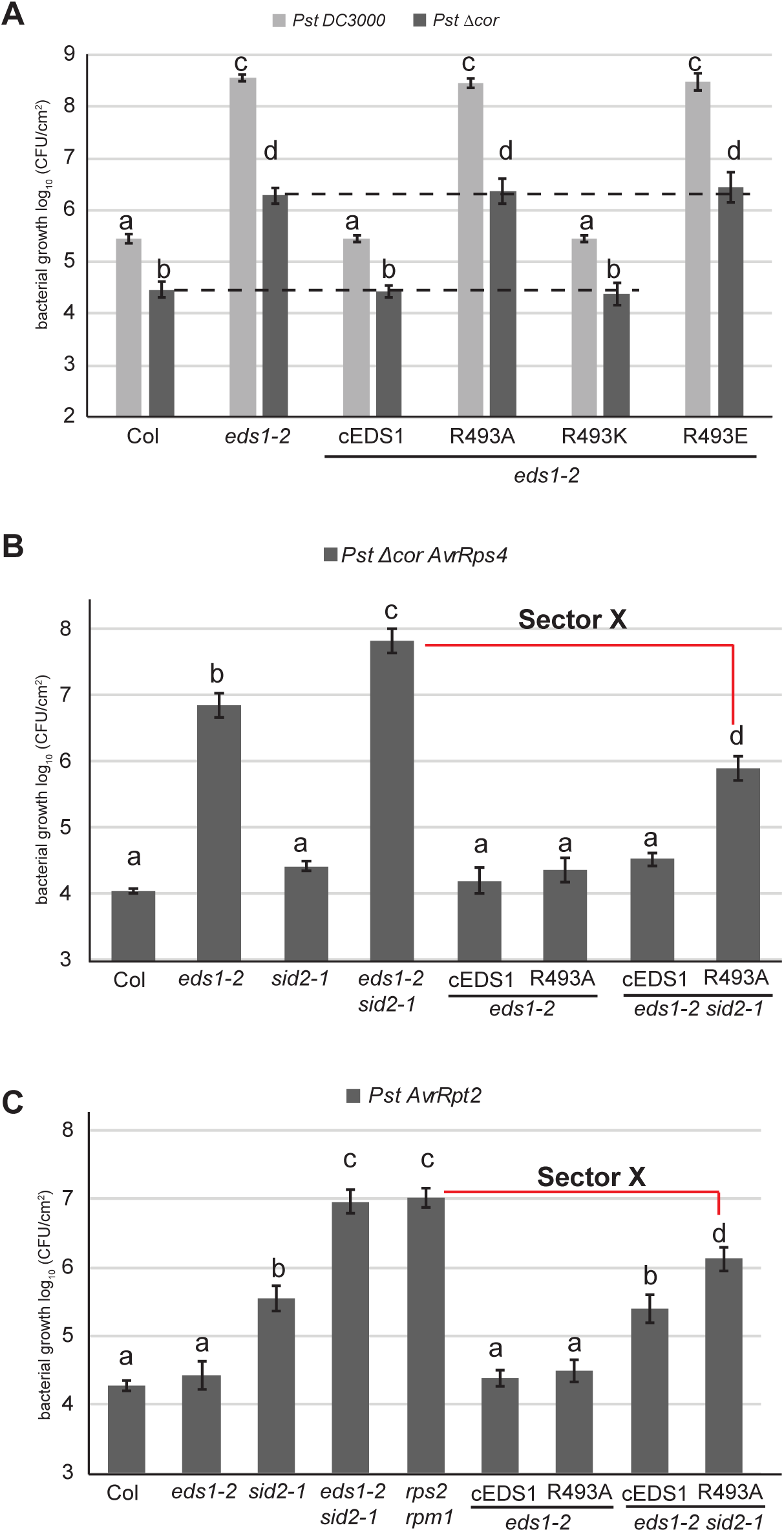
EDS1^R493^ signals in TNL and CNL (*RPS2*) immunity. **A**. Four-week-old Arabidopsis plants of the indicated genotypes were infiltrated with *Pst* DC3000 and *Pst* ΔCor (OD_600_ - 0.0005). Bacterial titers were determined at 0 and 3 dpi. No significant difference was observed between lines and treatments at 0 dpi. Bars represent mean of four biological replicates ± SE. Differences between genotypes were analysed using ANOVA (Tukey’s HSD, p<0.005). Similar results were obtained in three independent experiments. **B**. Four-week-old Arabidopsis plants of the indicated genotypes were infiltrated with *Pst* Δ*cor AvrRps4* (OD_600_ - 0.0005). Bacterial titers were determined at 0 and 3 dpi. No significant difference was observed at 0 dpi. Bars represent mean of four biological replicates ± SE. Differences between genotypes were analysed using ANOVA (Tukey’s HSD, p<0.005). Similar results were obtained in three independent experiments. C. Four-week-old Arabidopsis plants of the indicated genotypes were infiltrated with *Pst AvrRpt2* (OD_600_ - 0.0005). Bacterial titers were determined at 0 and 3 dpi. No significant difference was observed at 0 dpi. Bars represent mean of four biological replicates ± SE. Differences between genotypes were analysed using ANOVA (Tukey’s HSD, p<0.005). Similar results were obtained in three independent experiments.

### EDS1^493A^ defect is buffered by SA in TNL and CNL (*RPS2*) immunity

The *ICS1/SA* pathway can partially buffer *eds1* loss-of-function in TNL (*RRS1S RPS4*) ETI (Cui *et al*, 2018). Therefore, we tested the consequence of removing the ICS1/SA resistance sector and bacterial COR effects on R493A function in TNL ETI. For this, cEDS1 and R493A transgenic lines were crossed into an *eds1 ics1* (*eds1-2 sid2-1*) mutant background and growth of *Pst* Δ*cor AvrRps4* was measured at 3 dpi. As expected, both cEDS1 and R493A (in *eds1-2*) conferred full TNL immunity to *Pst* Δ*cor AvrRps4*, as did cEDS1 in *eds1-2 sid2-1* (Fig. 6B), consistent with EDS1 covering for loss of ICS1-generated SA in TNL ETI (Cui *et al*, 2017). By contrast, R493A *eds1-2 sid2-1* supported intermediate *Pst* Δ*cor AvrRps4* growth between cEDS1 *eds1-2 sid2-1* and *eds1-2 sid2-1* (Fig. 6B). Thus, *ICS1* generated SA compensates for R493A weak function in *RRS1S RPS4* ETI.

The ICS1/SA pathway compensates fully for loss of *EDS1* in CNL ETI mediated by two CNL receptors (*RPS2* and *HRT*(HR to turnip crinkle virus)) (Venugopal *et al*, 2009; Cui *et al*, 2017). We tested CNL (*RPS2*) immunity in the above R493A lines by infiltrating leaves with *Pst AvrRpt2*. In these bacterial growth assays, R493A in *eds1-2 sid2-1* also exhibited intermediate susceptibility between cEDS1 *eds1-2 sid2-1* and *eds1-2 sid2-1* (Fig. 6C). Together, these data show that the same EDS1 EP-domain surface functions in bacterial resistance mediated by a TNL receptor pair and a CNL receptor, and that in both ETI systems the weak resistance signalling activity of EDS1^R493A^ is buffered by *ICS1* generated SA. Interestingly, after stripping away SA and COR/JA effects, a portion of *EDS1* controlled resistance was detected in R493A *eds1-2 sid2-1* compared to *eds1-2 sid2-1* plants (Fig. 6B). Because the remaining resistance in R493A plants is independent of SA/COR effects in TNL ETI, we refer to it as sector X in a model (Fig.7).

**Figure 7.**
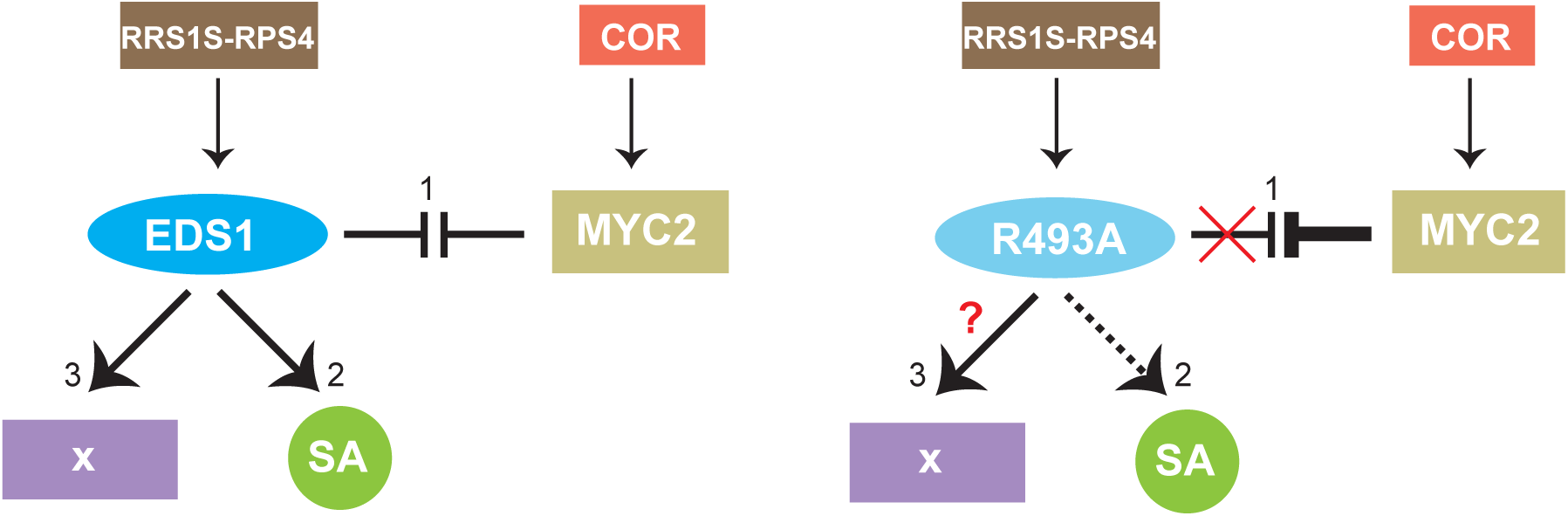
A model of EDS1 signalling branches in RRS1S RPS4 ETI. A three-pronged ETI signalling model derives from comparisons of wild-type EDS1, EDS1^R493A^ and *eds1-2* phenotypes in this study. Three interconnected EDS1 functions contribute to robust TNL ETI. 1) TNL-activated wild-type EDS1 effectively counters COR antagonism of immunity gene expression via MYC2. The defective EDS1 EP-domain mutant R493A is susceptible in TNL ETI against *Pst AvrRps4* due to its inability to counter COR/MYC2 antagonism. 2) EDS1 boosts SA accumulation independently of antagonising MYC2 while EDS1^R493A^ delays SA accumulation (dashed lines) independently of COR repressive effects. 3) An additional EDS1 branch (X) in TNL (*RRS1S RPS4*) ETI is revealed after removing ICS1/SA and COR effects. The EDS1 EP-domain, and more specifically EDS1^R493^, is also necessary for this resistance branch. The nature of branch X requires further study.

## Discussion

In Arabidopsis, EDS1-PAD4 and EDS1-SAG101 heterodimers, formed principally by the two partner N-terminal lipase-like domains, are required for TNL-triggered ETI and basal immunity against host-adapted bacterial (*Pst*) and oomycete (*Hpa*) pathogens (Parker *et al*, 1996; Feys *et al*, 2001; Feys *et al*, 2005; Jirage *et al*, 1999; Wagner *et al*, 2013). Current evidence suggests that the partner N-terminal lipase-like domains with characteristic α/β-hydrolase folds serve as a non-catalytic scaffold to stabilize the heterodimer, bringing together the essential α-helical EP-domains to form a cavity (Rietz *et al*, 2011; Wagner *et al*, 2013). The EP-domains have no known structural homologies outside the EDS1 family (Wagner *et al*, 2013). Here, using a combination of structure-guided mutants and reductionist genetic approaches, we identify conserved, positively charged residues (K440/441, K478 and R493) lining the EDS1 EP-domain cavity which are crucial for immunity signalling beyond heterodimer formation. By characterizing EDS1 partial‐ and loss-of-function mutations in the EP-domain, we establish that timely transcriptional mobilization of host immune response pathways is necessary for effective TNL ETI to *Pst* bacteria and that an important early EP-domain function of EDS1 complexes is to counter COR repression of immunity genes.

We focused our analysis on a single conserved, positively charged EP-domain residue, EDS1^R493^, because mutation of this to a neutral alanine (R493A) caused complete loss of TNL immunity to *Pst AvrRps4* and two *Hpa* strains, EMWA1 and CALA2 (Fig. 1B, C, E), indicating its broad importance for TNL ETI. A positive EDS1^R493^ character rather than the arginine *per se* is required for ETI activity because an R493K exchange behaved as wild-type EDS1, and R493E as the null *eds1-2* mutant, in *Pst AvrRps4* and *Hpa* EMWA1 infection assays (Fig. 5A, B). Thus, altering the charge of the amino acid side-chain at position R493 has a strong effect on heterodimer function without disturbing its interfaces (Fig. 5A, B, S4B). Maintained partner interactions of the EDS1^R493A^, EDS1^R493K^ and EDS1^R493E^ variants coupled with current molecular modelling suggest that the charge differences are unlikely to destroy integrity of the EP-domain but rather modify ionic interactions at the EP-domain cavity. The cavity surface around EDS1^R493^ is comprised of several conserved positively charged residues (Fig. S1A). A similar EP-domain surface was identified in PAD4 outside the cavity in which R420 forms comparable interactions with neighbouring glutamic acid and tyrosine residues as EDS1^R493^ (Fig. 1D). However, a PAD4^R420A^ exchange did not compromise TNL (*RPP4*) ETI (Fig. 1E), suggesting that the location and charge of EDS1^R493^ within the EP cavity are essential for EDS1-PAD4 function. We anticipate that functional interactions between the heterodimer EP-domains and other proteins or molecules are determined by the charge status of EDS1^R493^ and likely other positive residues (Fig. S1C). The JA/COR stimulated master TF MYC2 is not a candidate for direct EP cavity binding because MYC2 likely associates indirectly with EDS1-PAD4 complexes (Cui *et al*, 2018), and their association was not obviously affected by different mutations at EDS1^R493^ (Fig. S4C). Also, the partial ability of EDS1^R493A^ to antagonize COR/MYC2 controlled JA-branch genes at 24 hpi in *RRS1S RPS4* ETI to *Pst AvrRps4* did not correlate with altered EDS1-PAD4 interaction with MYC2 (Fig. S5B).

One striking feature of EDS1^R493A^ is that loss of TNL immunity to *Pst AvrRps4* is conditional on bacterial COR signalling via *MYC2*, as indicated by *Pst AvrRps4* and *Pst* Δ*cor AvrRps4* growth differences in R493A transgenic lines and fully restored resistance to *Pst AvrRps4* in an R493A *eds1-2 myc2-3* background (Fig. 3A, B). By contrast, EDS1^R493A^ behaves as a complete loss-of-function mutation in basal resistance to virulent *Pst DC3000* or *Pst* Δ*cor* (Fig. 6A). Thus, TNL (*RRS1S RPS4*) effector recognition and/or activation (Williams *et al*, 2014; Sarris *et al*, 2015; Le Roux *et al*, 2015) appears to equip EDS1 complexes, via the EP-domain positively charged surface, to block COR/MYC2 stimulated bacterial growth (Fig. 3A, 5B). We reported previously genetic and molecular evidence that TNL/EDS1 interference with COR stimulated MYC2 activity occurs inside nuclei coincident with or after COR is sensed by host COI1-JAZ complexes to release MYC2 from repression (Kazan & Manners, 2013; Zhang *et al*, 2015a; An *et al*, 2017; Cui *et al*, 2018). An additional TNL (*RRS1SRPS4*) ETI output might be to lower bacterial COR production during infection because an *in planta Pst* transcriptomic study showed that *Pst* COR biosynthesis genes were down-regulated at 6 hpi as part of the *RRS1S RPS4* immune response (Nobori *et al*, 2018).

Importantly, loss of TNL (*RPP4*) immunity to *Hpa* EMWA1 in R493A was not recovered by mutation of *MYC2* (Fig. 3C). Therefore, EDS1 heterodimers, via the EP-domains, must mobilize other anti-microbial pathways or processes independently of antagonizing COR/MYC2 signalling for TNL immunity against *Hpa*. It will be interesting to use the plant mutant combinations described here to pare down which pathways are responsible for stopping *Hpa* growth, building on earlier gene expression microrarray analyses of ETI responses to *Hpa* isolates (Eulgem *et al*, 2004; Wang *et al*, 2011). A number of protein effectors delivered to plant host cells by *Hpa*, fungal pathogens and *P. syringae* bacteria suppress SA immunity by targeting JA signalling to shift hormone balance away from SA (Kazan & Lyons, 2014; Yang *et al*, 2017). For example, *P. syringae* type III-secreted effectors HopX1 and HopZ1 derepress JA response genes independently of bacterial COR (Jiang *et al*, 2013; Gimenez-Ibanez *et al*, 2014). *P. syringae* effector HopBB1 activates a subset of JA outputs by increasing the binding between two JA response repressors, JAZ3 and TCP14, leading to their COI1-mediated elimination (Yang *et al*, 2017). Hence, microbes target SA-JA crosstalk in various ways to promote infection. Because EDS1 complexes effectively block bacterial COR virulence in *RRS1S RPS4* ETI (Fig. 3A, 5B) and in our RNA-seq data comparisons, we find a significant overlap between TNL/EDS1-dependent DEG and SA/JA-responsive genes (Fig. S3A, 4D), we speculate that one conserved role of EDS1 heterodimers is to protect the plant phytohormone network from interference by multiple pathogen effectors at various junctions, in order to preserve SA immunity.

A further insight to TNL/EDS1 ETI gained from our analysis is that perturbation of the EDS1 EP-domain in R493A lines causes an intrinsic delay in transcriptional reprogramming of Arabidopsis defence pathways, independently of bacterial COR. This is seen most clearly at the level of free (active) SA accumulation at 8 hpi and 24 hpi (Fig. 2A, 4A) and in RNA-seq analyses of wild-type Col (or cEDS1), *eds1-2* and R493A responses at 8 hpi and 24 hpi with *Pst AvrRps4* and *Pst* Δ*cor AvrRps4* (Fig. 4B, C). Therefore, timing of TNL/EDS1 induction of the *ICS1/SA* resistance branch and probably numerous other anti-microbial outputs (before or at 8 hpi) is not dictated by bacterial COR but the plant vulnerable to COR-producing *Pst AvrRps4* infection (Fig. 3A, 6B), and possibly also to TNL-recognized *Hpa* strains (Fig. 1B, E).

Partial recovery of gene expression changes in R493A lines at 24 hpi with *Pst AvrRps4* without restoration of *RRS1S RPS4* immunity (Fig. 2, 4B, S5B), emphasizes that there is a critical expression time-window for ETI to succeed. Based on these data, we propose that effective TNL/EDS1 immunity involves a crucial early step for rapid transcriptional activation of a broad set of defence pathways, rather than selectively mobilizing resistance outputs. This makes sense because various ETI and basal immune responses differ principally in timing and amplitude of transcriptional reprogramming, while the topology of co-expression networks appears to be quite stable (Tao *et al*, 2003; Tsuda *et al*, 2009; Kim *et al*, 2014; Hatsugai *et al*, 2017; Mine *et al*, 2018). Moreover, ETI specifically triggered by one pathogen effector or strain confers broad-spectrum immunity to a range of pathogen types (Rentel *et al*, 2008).

A recent time-resolved transcriptome study using single and higher order mutants of hormonal pathways indicates broad-sweep transcriptional changes rather than pathway specific signalling cascades in ETI (Mine *et al*, 2018), stressing the limited value of strong loss-of-function mutations in identifying causal resistance outputs. By comparing the *Pst AvrRps4* and *Pst* Δ*cor AvrRps4* transcriptomes in a weak EDS1^R493A^ mutant we were able to filter an interesting set of immune-related DEG (cluster #17) (Fig. 4C, Table S3). These genes would not have emerged from analysis of the cEDS1 and null *eds1-2* mutant responses alone because they were repressed at 8 hpi with *Pst AvrRps4* only in R493A tissues (Fig. 4C). Cluster #17 contains a number of functionally defined sensor *NLR* and *WRKY* TF genes, and one member (*ADR1-L2*) of a family of conserved helper NLRs (Fig. 4C) which contribute to ETI (Bonardi *et al*, 2011; Schön *et al*, 2013; Dong *et al*, 2016). Only one third of cluster #17 genes are represented in ‘ETI-related’ gene sets extracted from *RPS2* (CNL)-triggered transcriptomic analyses (Fig. S3B) (Hatsugai *et al*, 2017; Mine *et al*, 2018). The two thirds of cluster #17 ‘specific’ genes might represent another layer of immunity gene expression control in TNL ETI. It is tempting to speculate that susceptibility of R493A to *Hpa* (Fig. 1B, C) might be due in part to a failure to block *Hpa* effectors from targeting some or all of these genes for repression.

We have identified EDS1 EP-domain residues responsible for timely activation of transcriptional reprogramming, beyond heterodimer formation. Analysis of EDS1^R493A^ reinforces a two-pronged EDS1 transcriptional mechanism for *RRS1S RPS4* ETI proposed by Cui et al. (2018) which is (i) promotion of *ICS1* generated SA and (ii) blocking COR/MYC2 suppression of SA immunity. Delayed free SA accumulation in R493A plants at 8 hpi with *Pst AvrRps4* and *Pst* Δ*cor AvrRps4* (Fig. 4A) highlights a failure of this weak EDS1 allele to rapidly mobilize the SA-branch independently of COR. Early antagonism of COR/MYC2 signalling is important in *RRS1S RPS4* ETI because EDS1^R493A^ is fully susceptible to *Pst AvrRps4* despite being able to suppress COR/MYC2 marker genes almost as well as cEDS1 at 24 hpi (Fig. 2C, 4C, S5B). Notably, EDS1^R493A^ retains a portion of *RRS1S RPS4* (TNL) resistance to *Pst* Δ*cor AvrRps4* after removal of *ICS1* in an *eds1-2 sid2-1* double mutant (Fig. 6B). We therefore add a third yet unexplored EDS1 ETI signalling branch (denoted X in Fig. 7) which is independent of SA and JA/COR crosstalk. By testing R493A *eds1-2 sid2-1* plants in ETI conferred by a CNL receptor (*RPS2* recognizing *Pst AvrRpt2*) (Kunkel *et al*, 1993; Venugopal *et al*, 2009) (Fig. 6C), it becomes clear that the EDS1 EP-domain confers a crucial function in immunity triggered by TNL and certain CNL receptor types, which is worth exploring further.

## Materials and Methods

### Plant materials, growth conditions and pathogen strains

All mutants and lines are in the Col-0 genetic background. The mutants *eds1-2, sid2-1, eds1-2 sid2-1, myc2-3, pad4-1 sag101-3, eds1-2 myc2-3*, as well as YFP-cEDS1 and gEDS1-YFP *eds1-2* transgenic lines were previously described (Cui *et al*, 2018; Wagner *et al*, 2013; García *et al*, 2010). *Pseudomonas syringae* pv. *tomato* (*Pst*) strain DC3000, *Pst* Δ*cor, Pst* DC3000 *AvrRps4* (*Pst AvrRps4*), DC3000 *Δcor AvrRps4 (Pst* Δ*cor AvrRps4*) and DC3000 *AvrRpt2* (*Pst AvrRpt2*) are described (Cui *et al*, 2017). Plants were grown on soil in controlled environment chambers under a 10 h light regime (150-200 μE/m^2^s) at 22 °C and 60 % relative humidity.

### Pathogen infection assays

For bacterial growth assays, *Pst AvrRps4* or *Pst* Δ*cor AvrRps4* (OD_600_=0.0005) in 10mM MgCl2 were hand-infiltrated into leaves of four-week-old plants and bacterial titers measured at 4h post infiltration (day 0) and day 3 as described (Feys *et al*, 2005). Each biological replicate consists of three leaf disks from different plants and data shown in each experiment is compiled from 3-4 biological replicates. Statistical analysis was performed either using one-way ANOVA with multiple testing correction using Tukey’s HSD (p<0.005).

For gene expression and protein accumulation assays, leaves from four-week-old plants were hand-infiltrated with bacteria (OD_600_=0.005) and samples taken at indicated time points. For measuring protein accumulation, samples were pooled from at least three different plants. For gene expression analysis by qRT-PCR, four or more leaves from different plants were pooled as one biological replicate and two biological replicates were used in each independent experiment. Statistical analysis was performed by Student’s t-test with multiple testing correction using Bonferroni method (p<0.05).

*Hpa* isolates EMWA1 and CALA2 were sprayed on 2-3 week-old plants at 4*10^4^ spores/ml dH_2_O. Plant host cell death and *Hpa* infection structures were visualized under a light microscope after staining leaves with lactophenol trypan blue as described (Muskett *et al*, 2002). T_1_ complementation assays of Arabidopsis transgenic lines were performed as previously described (Stuttmann *et al*, 2011) and Hpa-infected seedlings rescued by spraying with Ridomil Gold (Syngenta). To quantify *Hpa* sporulation on leaves, three pots of each genotype were infected and treated as biological replicates. Plants were harvested at 6 dpi, their fresh weight determined, and conidiospores suspended in 5-10 ml dH_2_O and counted under the microscope using a Neubauer counting chamber.

### RNA isolation, library preparation and sequencing

For RNA-seq experiments, Col, *eds1-2*, cEDS1 and R493A#1 four-week-old plants were infiltrated with *Pst AvrRps4* or *Pst* Δ*cor AvrRps4* using the same bacterial titer as for gene expression assays. To randomize samples and reduce variation, total RNA was isolated from four individual plants per genotype (three infected leaves per plant) and pooled as one biological replicate. Each biological replicate is derived from an independent experiment. Total RNA was purified with an RNeasy Plant Mini Kit (Qiagen) according to manufacturer’s instructions. RNA-seq libraries were prepared from 1 μg total RNA according to TruSeq RNA sample preparation v2 guide (Illumina). Library construction and RNA sequencing was done by the Max-Planck Genome Centre (MPIPZ, Cologne), and produced 21-32 million 100 bp reads per sample. RNA-seq reads were mapped to the annotated genome of *Arabidopsis thaliana* (TAIR10) using TOPHAT2 (a = 10, g = 10) (Kim *et al*, 2013) and transformed into a read count per gene per sample using the htseq-count script (s = reverse, t = exon) in the package HTSeq (Anders et al., 2015). Genes with < 100 reads across samples were discarded. The count data from the remaining genes were TMM-normalized and log2 transformed using functions ‘calcNormFactors’ (R package EdgeR; (Robinson et al., 2009)) and ‘voom’ (R package limma; (Law et al., 2014)). To analyze differential gene expression over time between the different genotypes and treatments, for each analysis we fitted a linear model to the respective log_2_-transformed count data using the function lmFit (R package limma; (Law et al., 2014)) and subsequently performed moderated t-tests for specific comparisons of interest. In all cases, the resulting p values were adjusted for false discoveries due to multiple hypothesis testing via the Benjamini-Hochberg procedure. For each comparison, we extracted a set of significantly differentially expressed genes between the tested conditions (adjusted p-value ≤ 0.05, |log2FC| ≥ 1). RNA-seq experiments for *Pst AvrRps4* and *Pst* Δ*cor AvrRps4* were performed in separate batches and therefore normalized to Col-0 for the respective treatments to negate potential batch effects. The normalized values were used to generate a heatmap with hierarchial clustering. Circos plot was created using the R package ‘Circlize’ (Gu *et al*, 2014), to show the overlap of cluster #17 with other datasets.

### qRT-PCR analysis

Total RNA was extracted using a Plant RNA kit (Bio-budget). 500 ng total RNA was used for cDNA synthesis (quanta bio) and qRT-PCR analysis was performed using SYBR green master mix. The housekeeping gene *GapDH* was used as reference.

### Plasmid constructs

The pENTR/D-TOPO-cEDS1, pENTR/D-TOPO-gEDS1 and pENTR/D-TOPO cPAD4 vectors used for site-directed mutagenesis are previously described (Wagner *et al*, 2013; Stuttmann *et al*, 2016). Site-directed mutagenesis on the entry vectors was performed according to the QuikChange II site-directed mutagenesis manual (Agilent). Mutated entry clones were verified by sequencing and recombined into a pAM-PAT-based binary vector backbone by LR reaction.

### Generation of Arabidopsis transgenic plants

Stable transgenic lines were generated by transforming binary expression vectors into Arabidopsis null mutants *eds1-2* or *pad4-1 sag101-3*, as indicated, using *Agrobacterium-mediated* floral dipping.

### Yeast two-hybrid assays

Yeast 2-hybrid (Y2H) assays were performed using the Matchmaker system (Clontech) with strain AH109. Gateway cassettes were cloned into pGADT7 and pGBKT7 plasmids. Newly made PAD4 and EDS1 site-directed EP-domain mutants and the EDS1^LLIF^ variant (Wagner *et al*, 2013) were recombined into these plasmids by LR reaction. pGAD‐ and pGBK-containing co-transformants were selected on plates lacking leucine and tryptophan (-LW). Single colonies were re-streaked on plates additionally lacking histidine and adenine (-LWHA) to monitor reporter activation. Yeast growth was recorded after 2-5 d incubation at 30°C.

### Transient expression in Arabidopsis protoplasts

Leaf mesophyll protoplasts were prepared from 4-week-old *eds1-2 pad4-1 sag101-3* plants and transfections with plasmid DNA were done according to (Yoo et al., 2007). After transfection, protoplasts were incubated at room temperature under weak light (1.5 μE/m^2^s) for 16 h. Protoplasts were harvested and IPs performed as described below.

### Protein extraction, immunoprecipitation (IP) and immunoblotting

Total leaf extracts or protoplasts were processed in extraction buffer (50 mM Tris pH 7.5, 150 mM NaCl, 10 % (v/v) glycerol, 2 mM EDTA, 5 mM DTT, protease inhibitor (Roche, 1 tablet per 50 ml, 0.1 % Triton). Lysates were centrifuged for 15 min, 12000 rpm at 4 °C. 50 μl of supernatant was used as input sample. Immunoprecipitations (IPs) were conducted by incubating the input sample with 12 μl GFP-TrapA beads (Chromotek) for 2 h at 4°C. Beads were collected by centrifugation at 2000 rpm, 1 min at 4°C. Beads were washed three times in extraction buffer and boiled at 95°C in 2×Laemmli buffer for 10 min. Proteins were separated by SDS-PAGE and analyzed by immunoblotting. Antibodies used were α-GFP (Sigma Aldrich, 11814460001), α-HA (Sigma Aldrich, 11867423001), α-FLAG (Sigma Aldrich, F3165). Secondary antibodies coupled to Horseradish Peroxidase (HRP) were used for protein detection on blots (Santa Cruz Biotechnology and Sigma).

### SA quantitation

Free SA was quantified from leaf tissues (70-200 mg fresh weight), of four-week-old plants using a chloroform/methanol extraction and analysed by gas chromatography coupled to a mass spectrometer (GC-MS, Agilent), as described (Straus et al., 2010). Statistical analysis was performed by Student’s t-test with multiple testing correction using the Bonferroni method (p<0.05).

### Data Availability

The RNA-seq data are deposited in the National Center for Biotechnology Information Gene Expression Omnibus (GEO) database with accession number GSE116269.

## Acknowledgements

We thank Kenichi Tsuda and Takaki Maekawa (MPIPZ Cologne) for helpful discussions. This work was supported by The Max-Planck Society and Deutsche Forschungsgemeinschaft (DFG) grants within SFB 635 and SFB 670 (DDB), SFB 680 (DL), and an International Max-Planck Research School (IMPRS) doctoral fellowship (PvB).

## Author contributions

DDB and JEP designed the study; DDB and PVB performed experiments; BK, DDB and DL analysed the RNA-seq data; DDB and JB generated transgenic plant lines; KN provided structural insight; DDB and JEP wrote the paper with inputs from DL

## Supplementary figure legends

**Figure S1**

**A**. Crystal structure of EDS1 (blue) - SAG101 (green) heterodimer with cavity formed by the EP-domains (magenta mesh) highlighted. Conserved positively charged residues lining the cavity are depicted as brown sticks. **B**. Y2H interactions between activation domain (AD) fusions of EDS1 variants and PAD4 (BD) binding domain fusions. The EDS1-LLIF mutant which does not bind PAD4 was used as a negative control. Yeast viability (-LW) and protein interaction (-LWAH) are shown in the GAL4 matchmaker Y2H system. **C**. Summary of TNL (*RPP4*) complementation assay in T_1_ plants expressing EDS1-YFP EP-domain mutants. For each EDS1 mutant line, individual BASTA-resistant T_1_ seedlings were monitored for TNL-triggered resistance to *Hpa* EMWA1 (at 5 dpi). Seedlings showing conidospores on leaves were scored as disease susceptible. Numbers of resistant / total plants tested is shown. Non-transgenic controls for *RPP4* resistance Col-0 and *eds1-2* were not treated with BASTA. **D**. Conservation of EDS1 EP-domain residues across EDS1 orthologs. Sequences were aligned using MUSCLE (Edgar, 2004) and positions corresponding to K478 and R493 are highlighted, including the characteristic “EPLDIA” conserved motif of the EP-domain.

**Figure S2**

**A**. cEDS1-FLAG variants transiently co-expressed with cPAD4-YFP in *eds1-2 pad4-1 sag101-3* protoplasts (input) were immunoprecipitated (IP) using α-GFP beads. Co-immunoprecipitated FLAG-tagged cEDS1 variants were detected using α-Flag antibodies. EDS1-LLIF which (migrates higher) does not bind PAD4 was used as a negative control. **B**. Confocal images of four-week-old leaves of transgenic *eds1-2* lines expressing YFP-tagged EDS1 or R493A, showing nucleocytoplasmic localization. Images were taken at 24 hpi with *Pst AvrRps4*. Images are representative of >30 cells per line. Scale bar = 25μm. **C**. Four-week-old Arabidopsis plants of the indicated genotypes were infiltrated with *Pst AvrRps4* (OD_600_ - 0.0005) and bacterial titers determined at 0 and 3dpi. No significant difference was observed at 0 dpi. Bars represent mean of three biological replicates ± SE. Differences between genotypes were analysed using ANOVA (Tukey’s HSD, p<0.005). Similar results were obtained in three independent experiments. **D**. Accumulation of EDS1-YFP protein in mock and *Pst AvrRps4* (24 hpi) treated plants of cEDS1, gEDS1 and respective R493A mutants on immunoblots probed using α-GFP antibody.

**Figure S3**

**A**. Comparison of expression profiles of genes regulated by ICS1/SA/(Tsuda et al., 2013) and repressed by JA (Hickmann et al., 2017; selected clusters (#15, 18, 19, 20, 23, 24) were chosen based on differential expression profile between cEDS1 and *eds1-2*) with our study using *Pst AvrRps4* and *Pst* Δ*Cor AvrRps4* (ΔCor). Bar plots are coloured based on genoytpe with a light tone for the ΔCor data. Statistical differences between genotypes were analysed within treatment using the Kruskal-Nemenyi test (p<0.001) as indicated by different annotations. **B**. A Venn diagram presenting the overlap between genes in cluster #17 for AvrRps4 ETI at 8hpi, genes that are specifically regulated in *AvrRpt2-*-triggered ETI (ETI-specific genes) (Mine et al., 2018), DEG at 4 hpi with *Pst AvrRpt2* (Pst AvrRpt2 - 4 hpi) (Mine et al., 2018) and DEG at 5 h with estradiol-induced AvrRpt2 (ED-AvrRpt2-5hpi) (Hatsugai et al., 2017).

**Figure S4**

**A**. Confocal images of transiently expressed FLAG-tagged cEDS1 and R493 mutant variants in *N. benthamiana* showing nucleocytoplasmic localization. White arrowheads depict nuclei and chloroplasts fluoresce red. Images are representative of >20 cells/variant at 3d after Agroinfiltration. Scale bar = 20μm. **B**. EDS1-FLAG variants transiently expressed with PAD4-YFP in *eds1-2 pad4-1* protoplasts were immunoprecipitated using α-GFP beads. Co-immunoprecipitated EDS1 variants were detected using α-FLAG antibodies. EDS1^LLIF^-FLAG and YFP-FLAG were used as negative controls in the IP. **C**. EDS1-FLAG variants, as used in B. were transiently expressed with PAD4-YFP and strep-HA-MYC2 (SH-MYC2) in *eds1-2 pad4-1 sag101-3* protoplasts. PAD4-YFP was immunoprecipitated using α-GFP beads. Co-immunoprecipitated FLAG-tagged EDS1 variants and SH-MYC2 were detected on immunoblots using α-FLAG and α-HA antibodies, respectively. EDS1^LLIF^-FLAG and YFP-FLAG were used as negative controls. IPs were repeated three times with similar results.

**Figure S5**

A. Expression of *EDS1*, *PAD4* and *ICS1* in EDS1 transgenic lines, as indicated, at 8 hpi with *Pst AvrRps4* or *Pst* Δ*cor AvrRps4*, measured by qRT-PCR. Values were normalized to the house-keeping gene *GapDH*. Bars represent means ± SE calculated from three independent experiments. Differences between genotypes were calculated using ANOVA (Tukey’s HSD, p<0.05). B. Expression of selected MYC2-marker genes *BSMT1*, *JAZ10* and *VSP1* in EDS1 mutant lines at 24 hpi with *Pst AvrRps4*, measured by qRT-PCR. Values were normalized to the house-keeping gene *GapDH*. Bars represent means ± SE calculated from three independent experiments. Differences between genotypes was calculated using ANOVA (Tukey’s HSD, p<0.05).

## Supplementary data tables

**Table S1**. List of DEG between R493A and *eds1-2* at 8 hpi with *Pst AvrRps4*.

**Table S2**. Summary of total DEG between *Pst AvrRps4* and *Pst* Δ*Cor AvrRps4* treatments for each genotype at 8 hpi and 24 hpi.

**Table S3**. List of genes in cluster #17 (associated with Fig. 4C, D).

**Table S4**. Gene Ontology (GO) enrichment of cluster #17 genes. GO enrichment was obtained using the panther database.

**Table S5**. Overlapping genes between cluster #17, BTH-regulated genes and JA-regulated genes (associated with Fig. 4D).

**Table S6**. List of primers used in this study.

